# Ensemble deep learning of embeddings for clustering multimodal single-cell omics data

**DOI:** 10.1101/2023.02.22.529627

**Authors:** Lijia Yu, Chunlei Liu, Jean Yee Hwa Yang, Pengyi Yang

**Affiliations:** Computational Systems Biology Group, Children’s Medical Research Institute, University of Sydney, Westmead, NSW 2145, Australia; School of Mathematics and Statistics, University of Sydney, NSW 2006, Australia; Charles Perkins Centre, University of Sydney, NSW 2006, Australia; Sydney Precision Data Science Centre, University of Sydney, NSW 2006, Australia; Laboratory of Data Discovery for Health Limited (D^2^4H), Science Park, Hong Kong SAR, China

## Abstract

**Motivation:** Recent advances in multimodal single-cell omics technologies enable multiple modalities of molecular attributes, such as gene expression, chromatin accessibility, and protein abundance, to be profiled simultaneously at a global level in individual cells. While the increasing availability of multiple data modalities is expected to provide a more accurate clustering and characterisation of cells, the development of computational methods that are capable of extracting information embedded across data modalities is still in its infancy.

**Results:** We propose SnapCCESS for clustering cells by integrating data modalities in multimodal singlecell omics data using an unsupervised ensemble deep learning framework. By creating snapshots of embeddings of multimodality using variational autoencoders, SnapCCESS can be coupled with various clustering algorithms for generating consensus clustering of cells. We applied SnapCCESS with several clustering algorithms to various datasets generated from popular multimodal single-cell omics technologies. Our results demonstrate that SnapCCESS is effective and more efficient than conventional ensemble deep learning-based clustering methods and outperforms other state-of-the-art multimodal embedding generation methods in integrating data modalities for clustering cells. The improved clustering of cells from SnapCCESS will pave the way for more accurate characterisation of cell identity and types, an essential step for various downstream analyses of multimodal single-cell omics data.

**Availability and implementation:** SnapCCESS is implemented as a Python package and is freely available from https://github.com/yulijia/SnapCCESS.

## 1 Introduction

The development of novel single-cell technologies, such as cellular indexing of transcriptomes and epitopes by sequencing (CITE-seq) (Stoeckius *et al*., 2017), shared single-cell profiling of RNA and chromatin (SHARE-seq) (Ma *et al*., 2020) and trimodal single-cell profiling by TEA-seq (Swanson *et al*., 2021), enables the profiling of gene expression, protein abundance, and/or chromatin accessibility in the same cell. The availability of multiple data modalities in individual cells promises more precise characterisation of cells such as clustering cells into distinctive cell types (Zhu *et al*., 2020). To analyse such multimodal single-cell omics data, however, requires effective computational methods that are capable of integrating data modalities for extracting the underlying biological signals.

While various methods exist for enabling the clustering of multimodal single-cell omics data, such as by simple feature concatenation or more sophisticated methods that integrate clustering output from each modality (Miao *et al*., 2021; Adossa *et al*., 2021), a popular approach is to integrate data modalities through learning an embedding that encodes multiple data modalities into a shared latent space, from which any clustering algorithm that accepts the embedding as input could be used for clustering cells (Lin *et al*., 2022). For example, Jvis-learn performs joint dimension reduction of data modalities for generating embeddings of multimodal single-cell omics data (Do and Canzar, 2021). It automatically determines the relative importance of each data modality that emphasises distinguishing characteristics while reducing noise. MOFA+ implements a Bayesian group factor analysis framework to infer a lowdimensional embedding that captures shared variation across multiple modalities (Argelaguet *et al*., 2020). Recently, deep learning-based methods such as totalVI (Gayoso *et al*., 2021) use a variational autoencoder (VAE) to learn an embedding for integrating RNA and ADT modalities such as in CITE-seq data, and MultiVI (Ashuach *et al*., 2021) uses a VAE to integrate RNA and ATAC modalities such as in SHARE-seq data. The multimodality integrated embeddings generated from these methods can be subsequently applied for cell clustering using any clustering algorithms that accept embeddings as input. Thus, the utility and quality of the multimodality integrated embeddings will have a large impact on clustering multimodal single-cell omics data.

Here we aim to improve the clustering of multimodal single-cell omics data by developing an ensemble deep learning-based framework that generates a multi-view of multimodality integrated embeddings. This is motivated by our previous work on autoencoder-based ensemble clustering of scRNA-seq data that demonstrates consensus derived from multiple embeddings each generated from perturbing the input data can lead to significantly better clustering results (Geddes *et al*., 2019). Such a single-cell consensus clusters of encoded subspaces (scCCESS) approach benefits from the multi-view of the input data (Cao *et al*., 2020) and is generic to clustering algorithms. Built on this concept, we propose SnapCCESS, an ensemble clustering framework that uses VAE and the snapshot ensemble learning technique (Huang *et al*., 2017) to learn multiple embeddings each encoding multiple data modalities, and subsequently generate consensus clusters for multimodal single-cell omics data by combining clusters from each embedding. The innovation in SnapCCESS includes (i) implementing an ensemble deep learning framework for creating a multi-view of latent spaces from which multimodality embeddings of multimodal single-cell omics data can be generated, and (ii) designing a snapshot ensemble learning approach to significantly improve the computational efficiency of the proposed framework.

By applying SnapCCESS to multimodal single-cell omics datasets generated by various biotechnologies and protocols, we show that SnapCCESS is effective and computationally efficient in learning multiple embeddings compared to conventional ensemble deep learning methods. We found that multimodal data generally offer more information than single modality alone and SnapCCESS leverages such information to improve cell clustering. We also show that embeddings learned from SnapCCESS are generalisable and can be coupled with various clustering algorithms for improving consensus clustering of multimodal single-cell omics data. Lastly, we demonstrate the competitive performance of SnapCCESS with the other state-of-the-art methods for generating embeddings of data modalities for clustering cells. Together, our work showcases the effectiveness of a novel unsupervised ensemble deep learning framework for performing clustering analysis of multimodal single-cell omics data.

## 2 Materials and methods

### 2.1 SnapCCESS framework for generating embeddings of multimodal single-cell data

To integrate the high-dimensional feature space in each modality of multimodal single-cell omics data, SnapCCESS encodes features from multiple data modalities into a latent space using a VAE model by jointly learning to reconstruct each data modality (**Figure 1a**).

**Figure 1.**
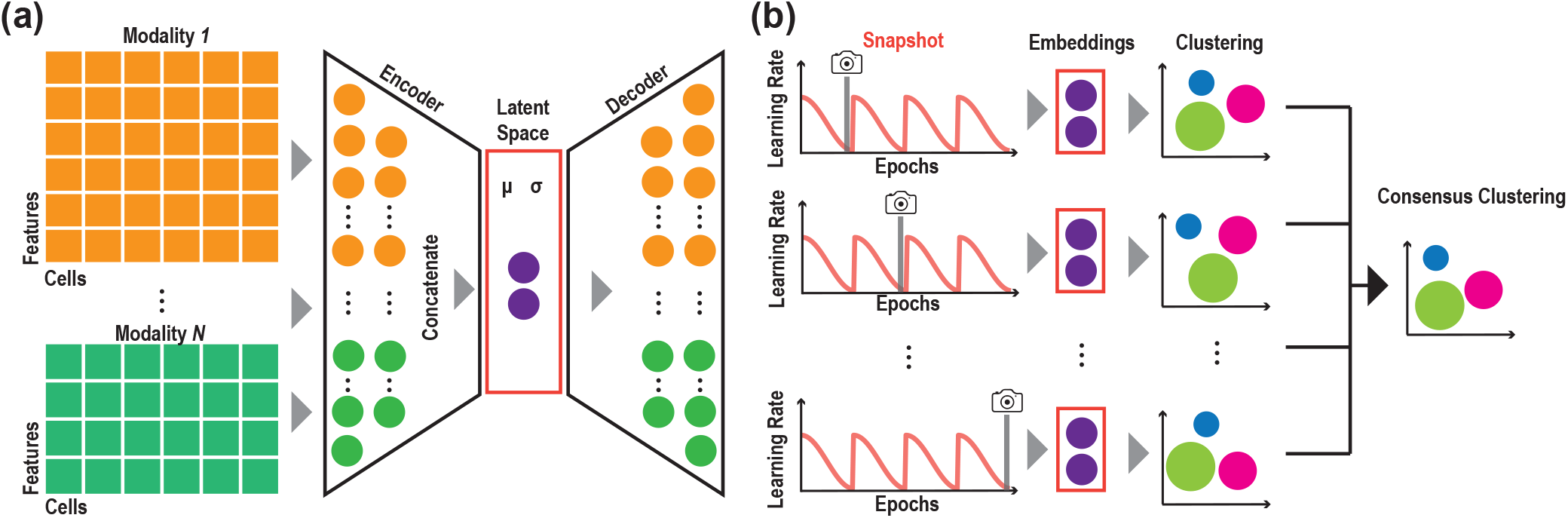
The proposed SnapCCESS framework of ensemble deep learning of embeddings for multimodal single-cell data clustering. (**a**) A variational autoencoder (VAE) is used to encode the highdimensional features from multimodal data to a low-dimensional latent space. (**b**) The training process of SnapCCESS is based on the snapshot ensemble deep-learning model using learning rate annealing cycles where the model converges to and then escapes from multiple local minima, and multiple snapshots were taken at these minima for creating a multi-view of embeddings. The schematic illustrates using epoch of 1 for generating snapshots. The consensus clustering result is derived from combining individual clustering results each from a snapshot embedding.

Specifically, SnapCCESS consists of multimodality-specific encoders and decoders for data integration and dimension reduction. The encoders in the VAE component include one learnable point-wise parameters layer and one fully connected layer to the input layer. Because surface protein modality has significantly fewer features than RNA and ATAC modalities, we empirically set the numbers of neurons for encoders of RNA, surface proteins, and ATAC modalities to be 185, 30, and 185, respectively. To learn a latent space that integrates the information across modalities, we concatenate the output from the encoder trained from each data modality to perform joint learning using a fully connected layer with 100 neurons. The embeddings of input data were obtained from the latent space of VAE by minimising the loss function *L* defined as follows:

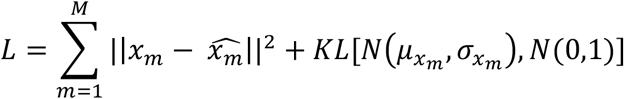

where *x* represents the original input, 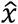 represents the reconstructed data, *i* is the *i^th^* modality, *M* is the number of modalities in a multimodal data. *N*(·) presents a normal distribution, which is learnable in VAE.

To implement the snapshot ensemble technique (Huang *et al*., 2017), the learning rate of the training model was set up to be the shifted cosine function, which could help the training model converge to multiple local minima and then get multiple lower-dimensional embeddings (**Figure 1b**). This is defined as follows:

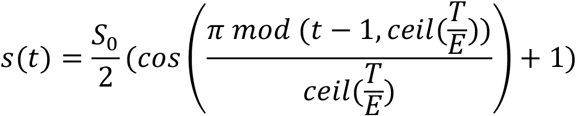

where *s*_0_ is the initial learning rate, *t* is the iteration number, *T* is the total number of training iterations, *E* is the number of learning rate cycles, *mod*(·) refers to the modulo operator.

### 2.2 Clustering algorithms for ensemble clustering of embeddings

To create consensus clustering, embeddings generated from SnapCESS and conventional VAE ensemble method were used for generating clustering results and then combined for deriving consensus results. Since the embeddings generated by these methods can be coupled with various clustering algorithms for creating consensus clustering results, we have included three different clustering algorithms for testing their effectiveness. These include a simple k-means clustering algorithm, a more sophisticated spectral clustering method, and SIMLR, a kernel-based clustering method designed for scRNA-seq data analysis (Wang *et al*., 2017). In particular, for the simple k-means clustering, we utilised the ‘kmeans’ function in the stats package with the default settings. For the spectral clustering algorithm, we employed the ‘spectralClustering’ function from the CiteFuse package (Kim *et al*., 2020) with the default parameters to perform spectral clustering. Lastly, we used the ‘SIMLR_Large_Scale’ function in the SIMLR package with the number of principal components set to 20 as recommended. For all clustering algorithms, the number of clusters was set to be the same as the number of cell types in each dataset based on the cell type annotation from the original study. After obtaining individual clustering output from each clustering method, a fixed-point iteration algorithm *g*(·) for obtaining hard least squares Euclidean consensus partitions was applied to compute the consensus clusters of individual partitions using clue package ‘cl_consensus’ function (Hornik, 2005):

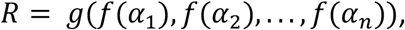

where *R* is the clustering result and *f*(·) represents performing a given clustering approach using an embedding *α_i_*. All clustering analyses were carried out in the R programming environment.

### 2.3 Evaluation settings and datasets

#### 2.3.1 Conventional VAE ensemble

Conventional ensemble deep learning typically relies on perturbing the initialisation and/or input data to train individual base models that can be used for creating the final ensemble model (Cao *et al*., 2020). To compare the performance of SnapCCESS with the conventional VAE ensemble method, we implemented a conventional VAE ensemble learning model by using random initialisation of the VAE neural networks. For a fair comparison, we used the same VAE model as in SnapCCESS to learn the latent space for integrating the data modalities in multimodal single-cell omics data. The same numbers of neurons for encoders and decoders were used as in SnapCCESS and the learning rate was fixed at 0.02. To obtain multiple embeddings, multiple VAEs were trained each with a different set of network initialisation weights on the input dataset.

#### 2.3.2 Settings for other multimodal embedding generation methods

##### MOFA+

MOFA2 (v1.6.0) which implements the MOFA+ algorithm (Argelaguet *et al*., 2020) was used to generate multi-modality embeddings of the six datasets. In accordance with the author’s tutorial (https://raw.githack.com/bioFAM/MOFA2_tutorials/master/R_tutorials/getting_started_R.html), prenormalised datasets were first used to create the mofa object using ‘create_mofa’ function. The data were then used as input for the modal training using ‘run_mofa’ function with default parameters. We use the ‘get factors’ function with factors = ‘all’ to obtain the embeddings for each input dataset.

##### Jvis-learn

Jvis-learn (v0.0.12) (Do and Canzar, 2021) was employed for generating multi-modality embeddings of the six datasets. The pre-normalised datasets were used as input for creating the j-SNE embeddings that joint multimodal omics data via the ‘JTSNE’ function.

##### totalVI

totalVI is designed for anaysing CITE-seq data (Gayoso *et al*., 2021). In this study, the totalVI procedure implemented in the scvi-tools package (v0.17.3) was used for generating multi-modality embeddings of the three CITE-seq datasets. Following the author’s tutorial (https://docs.scvi-tools.org/en/stable/tutorials/notebooks/totalVI.html), the raw count matrices of RNA and ADT were first normalised using the ‘normalize_total’ and ‘log1p’ functions and then top 4,000 most variable genes were selected using the ‘highly_variable_genes’ function. The data were subsequently used as input for model training using ‘scvi.model.TOTALVI.setup_anndata’, ‘scvi.model.TOTALVI’, and ‘train’ functions in scvi-tools. The latent space of RNA and ADT modalities was generated using the ‘get_latent_representation’ function.

##### MultiVI

MultiVI is a sibling of totalVI and specifically designed for anaysing data with RNA and ATAC modalites (Ashuach *et al*., 2021). Here, the MultiVI procedure implemented in the scvi-tools (v0.17.3) was used for multi-modality integration of the SHARE-seq and SNARE-seq datasets. In accordance with the author’s tutorial (https://docs.scvi-tools.org/en/stable/tutorials/notebooks/MultiVI_tutorial.html), the raw count matrices of RNA and gene activity score matrices from ATAC and the paired matrix of RNA and ATAC were utilised as input. These data were first concatenated using the ‘organize_multiome_anndatas’ function in scvi-tools and then used for model training using ‘scvi.model.MULTIVI.setup_anndata’, ‘scvi.model.MULTIVI’ and ‘train’ functions in scvi-tools. The latent space of RNA and ATAC modalities was generated using the ‘get_latent_representation’ function.

#### 2.3.3 Datasets and pre-processing

A collection of six multimodal single-cell omics datasets generated by different biotechnologies were included in this study for method evaluation. These include three CITE-seq datasets, one SNARE-seq dataset, one SHARE-seq dataset, and one TEA-seq dataset. For datasets with multiple samples and batches, one representative sample or batch was selected for analyses to avoid batch effects. Each dataset was filtered before the raw counts were transformed into log normalised counts using the ‘logNormCounts’ function in the scater package (McCarthy *et al*., 2017). The filtering process for each dataset is as below.

*Ramaswamy CITE-seq dataset* (Ramaswamy *et al*., 2021). The raw RNA and ADT matrices of PBMC from three healthy donors were downloaded from NCBI GEO using the accession number GSE166489. We used the healthy donor (GSM5073072) in our analysis. After filtering RNA and ADT expressed in less than 1% of the cells and genes, discarding cell types with fewer than 50 cells, we obtained 9,745 cells and 21 cell types, with 11039 RNA and 189 ADT features.

*Stephenson CITE-seq dataset* (Stephenson *et al*., 2021). The PBMC CITE-seq data of healthy individuals sequenced by NCL medical centre was used in this study. The raw matrices of RNA and ADT and the annotation of cells to their respective cell types from the original study were downloaded from the EMBL-EBI ArrayExpress database under the accession number E-MTAB-10026. RNA and ADT in this dataset were filtered by removing those that expressed in less than 1% of the cells and genes, cell types were filtered by removing those that have less than 50 cells. After filtering, 64,197 cells from 15 cell types (4999 RNA, 192 ADT) were kept for analysis.

*Hao CITE-seq dataset* (Hao *et al*., 2021). The raw RNA and ADT matrices from this CITE-seq dataset generated by Hao et al. from PBMC were downloaded from NCBI GEO under the accession number GSE164378. As the above, RNA and ADT in this dataset were filtered by removing those that expressed in less than 1% of the cells and genes, and cell types were filtered by removing those that have less than 50 cells. In total, 67,035 cells (11,451RNA, 228 ADT) and 29 cell types in batch 1 of the dataset were used in the analysis.

*Chen SNARE-seq dataset* (Chen *et al*., 2019). The SNARE-seq data that measures RNA and ATAC from matched cells in the adult mouse brain cortex sample (AdBrainCortex) was downloaded from NCBI GEO under the accession number GSE126074. The cell type information was obtained from the authors. For ATAC data, peaks with no expression across cells were removed. We then summarised the ATAC data from peak level into gene activity scores using the ‘CreateGeneActivityMatrix’ function in Seurat. We filtered out RNA and ATAC quantified in fewer than 1% of the cells and genes, and removed cell types that have less than 50 cells, resulting in a dataset with 9,930 cells (11,011 RNA, 16,443 ATAC peak features) and 20 cell types for the subsequent analyses.

*Ma SHARE-seq dataset* (Ma *et al*., 2020). The SHARE-seq data that measures RNA and ATAC from matched cells in mouse skin samples were downloaded from NCBI GEO under the accession number GSE140203. Similar to the above, we first removed peaks with no expression across cells, and then summarised the ATAC data from peak level into gene activity scores using the ‘CreateGeneActivityMatrix’ function in Seurat. We filtered out RNA and ATAC quantified in fewer than 1% of the cells and genes, and remove cell types that have less than 50 cells, resulting in a dataset with 32,968 cells (8,765 RNA, 17,413 ATAC peak features) and 23 cell types for the subsequent analyses.

*Swanson TEA-seq dataset* (Swanson *et al*., 2021). TEA-seq enables simultaneous single-cell profiling of transcripts, epitopes, and chromatin accessibility. The processed matrices of TEA-seq data from measuring PBMC were downloaded from the NCBI Gene Expression Omnibus (GEO) under the accession number GSE158013, with raw RNA expression, ADT expression, and peak accessibility (ATAC) measured for the same cells in four data batches. Due to the low batch effect presented in the four datasets, we merged the four data batches. We summarised the matrix of ATAC from peak level to gene activity scores using the ‘CreateGeneActivityMatrix’ function in the Seurat package. Genes with fewer than 1% quantifications across all cells and all genes in the three modalities are removed. This resulted in a dataset with 25,286 cells and 9 cell types, including 9,772 RNA, 46 ADT and 16,520 ATAC peak features.

## 2.4 Performance evaluation criteria

### Clustering concordance performance evaluation

For evaluating clustering performance, we used adjusted Rand index (ARI) and normalised mutual information (NMI) to evaluate the clustering concordance with respect to pre-defined cell type annotations from their original studies (Kim *et al*., 2019). Let S be a set of *N* cells, then a clustering *U* on *S* is a way of partitioning *S* into non-overlap subset {*U*_1_, *U*_2_,…, *U_R_*}. Here, we define *U* = {*U*_1_, *U*_2_,…, *U_R_*} as the real cell type labels with *R* cell types, *V* = {*V*_1_, *V*_2_,…, *V_c_*} is a partition with *C* clusters generated by a clustering. Pair counting based measures can be used for counting pairs of items on which the partition *U* and *V* agree or disagree. Specifically, the 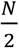. item pairs in *S* can be classified into one of the four types: (i) *N*_11_: the number of pairs that are in the same partition in both *U* and *V;* (ii) *N*_00_: the number of pairs that are in different partitions in both *U* and *V;* (iii) *N*_01_: the number of pairs that are in the same partition in *U* but in different partitions in *V*; (iv) *N*_10_: the number of pairs that are in different partitions in *U* but in the same partition in *V*. Following this, ARI, NMI can be defined as follows:

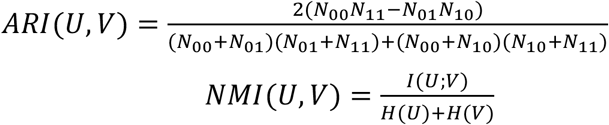

where *I*(*U*; *V*) is the mutual information between *U* and *V*, defined as:

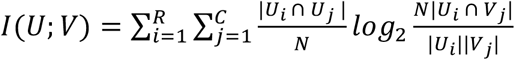

and *H*(·) is the entropy of partitions, in which *H*(*U*) and *H*(*V*) are calculated as:

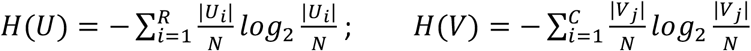

### Assessment of the run time usage

To evaluate the computation speed of SnapCCESS and the conventional VAE ensemble clustering method, all benchmark tasks were allocated to a research server with an NVIDIA GPU GeForce RTX 2080 Ti. The elapsed run time was calculated by using the Python function ‘time.perf_counter()’. Time for each method only takes into account the deep learning network building and model training steps.

## 3 Results

### 3.1 SnapCCESS is an effective and efficient ensemble deep learning method for clustering multimodal single-cell omics data

We first evaluated the performance of SnapCCESS and compared its performance with a conventional VAE ensemble clustering method on the six multimodal single-cell omics datasets generated from different biotechnological platforms. The concordance of the k-means clustering output from each method with respect to the cell type annotation from the original study of each dataset was quantified using ARI (**Figure 2a**) and NMI (**Figure 2b**) and the procedure was repeated 20 times to account for the variability in the clustering results. Notably, we found that ensemble learning improves the clustering performance of both methods and the clustering concordance increases while the variability reduces with the ensemble size (i.e. the number of base clusters). We also found that in general the improvement plateau at around the ensemble size of 50.

**Figure 2.**
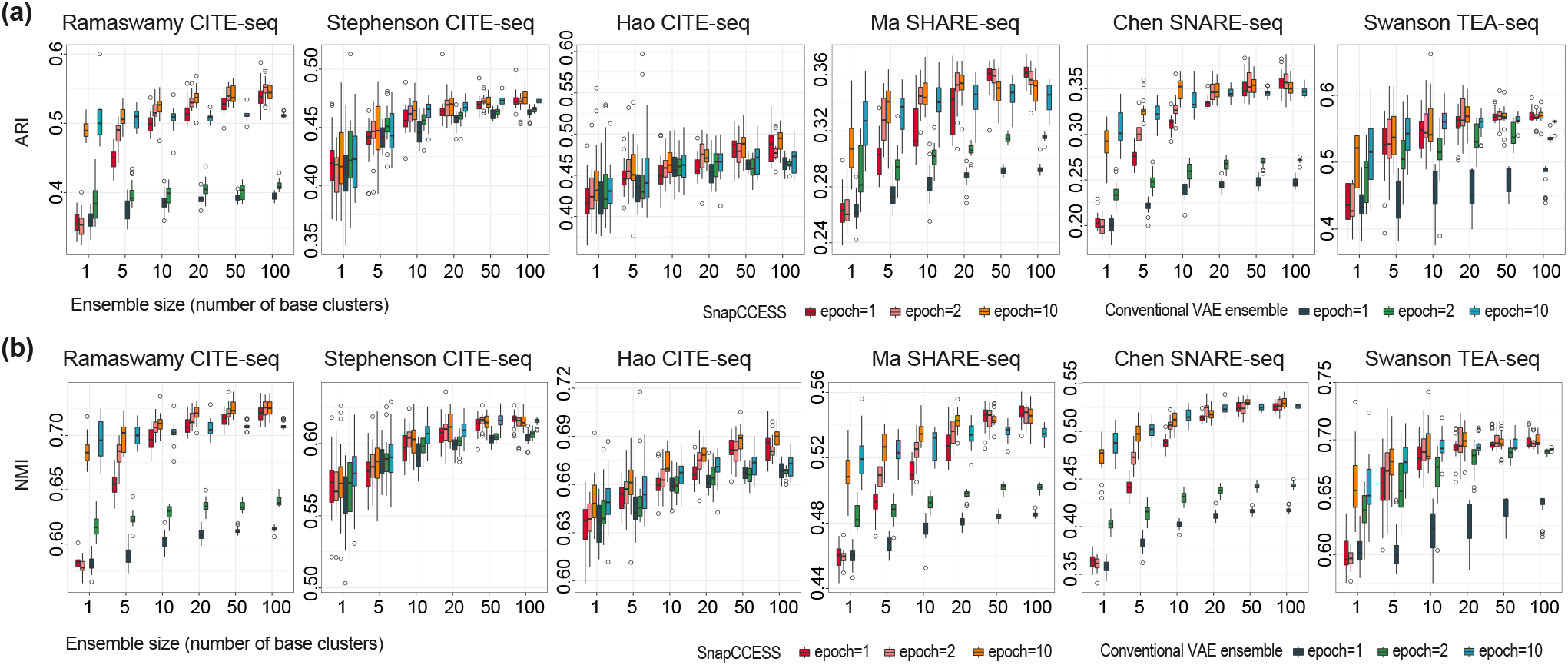
Clustering performance of SnapCCESS and conventional cluster ensembles trained by different numbers of epochs. (**a**) Concordance of cell type clustering on six multimodal single-cell omics data. The x-axis is the number of base clusters for the ensemble and the y-axis is the concordance of the clustering output and the cell type annotation in the original studies quantified by ARI. (**b**) Similar to (a) but with the clustering concordance quantified by NMI. The k-means clustering algorithm was used for clustering the embeddings generated from each method. The entire procedure was repeated 20 times for capturing the performance variability.

A key advantage of SnapCCESS is its ability to generate informative embeddings from multiple local minima and therefore requires much fewer epochs during the ensemble learning process (**Figure 1b**). To validate this, we tested using epochs of 1, 2, and 10 in SnapCESS and the conventional VAE ensemble clustering. As expected, in most cases the conventional VAE ensemble clustering requires a high number of epochs to achieve high performance (**Figure 2**). In comparison, SnapCESS achieves comparable performance using only one epoch in training the ensemble model at the sizes of 50 and 100. Since the epoch is a key parameter that defines the number of times the VAE will work through an input dataset, in general training on more epochs requires more computing time. To evaluate this, we recorded the computation time of SnapCESS and the conventional VAE ensemble on each dataset. Indeed, we found that, for both methods, fewer epochs resulted in significantly faster computation, especially with large ensemble sizes (**Figure 3**). Under the same number of epochs, the computation time of both methods is very similar. Nevertheless, from our above analyses, only SnapCCESS could achieve high performance in clustering cells with a low training epoch and significantly outperforms the conventional VAE ensemble with an epoch of 1. Taken together, these findings demonstrate that combining individual clustering results derived from multiple embeddings does lead to more accurate and reproducible consensus clustering of multimodal single-cell omics data, and the SnapCESS framework for ensemble deep learning of embeddings can achieve high-performance cell clustering using significantly less computational time.

**Figure 3.**
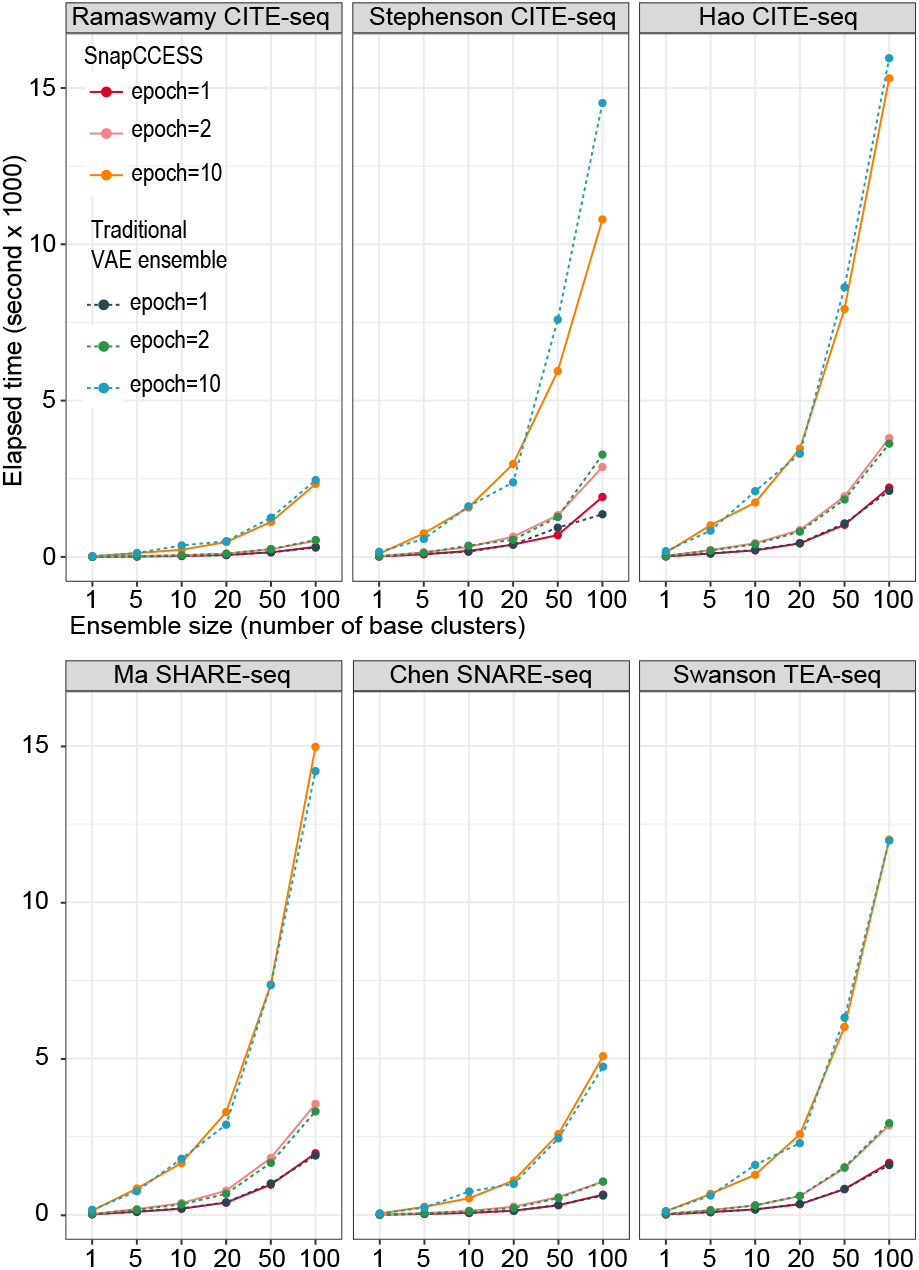
Comparison of computation time for SnapCCESS and conventional VAE cluster ensembles. The x-axis represents the number of base clusters included in the cluster ensembles and the y-axis is the elapsed time in the unit second. Results from SnapCCESS are presented as solid lines and those from conventional VAE cluster ensembles are presented as dashed lines. Epochs of 1, 2, and 10 were tested and denoted using different colours.

### 3.2 Integrated embedding of multimodality generally leads to more precise cell clustering compared to unimodality embeddings

One of the key motivations in conducting multimodal single-cell omics experiments is the anticipation that the availability of multiple molecular features in individual cells will lead to more precise characterisation of cell identity and heterogeneity in complex multicellular organisms and biological systems (Zhu *et al*., 2020). To investigate this, for each dataset, we trained SnapCCESS (epoch=1) using either all available data modalities or each unimodality independently, and then performed cell clustering using the k-means clustering algorithm on either the integrated embedding of multimodality or embeddings from each unimodality. We found that in general clustering of cells using integrated embeddings of multimodality leads to significantly better results than from using any unimodality alone (**Figure 4**). Among the clustering results using unimodal embeddings, those generated from RNA modality generally performed similarly or better compared to those from ADT modality. The clustering performance of ATAC modality appears to be lower compared to other modalities, which may be due to the higher dimensionality and data sparsity in ATAC data modality (Xiong *et al*., 2019). Together, these results support the expectation that taking into consideration of multiple molecular features of cells can lead to a more precise downstream characterisation of the biological systems, and further highlight the utility of modality integration methods for analysing multimodal single-cell omics data.

**Figure 4.**
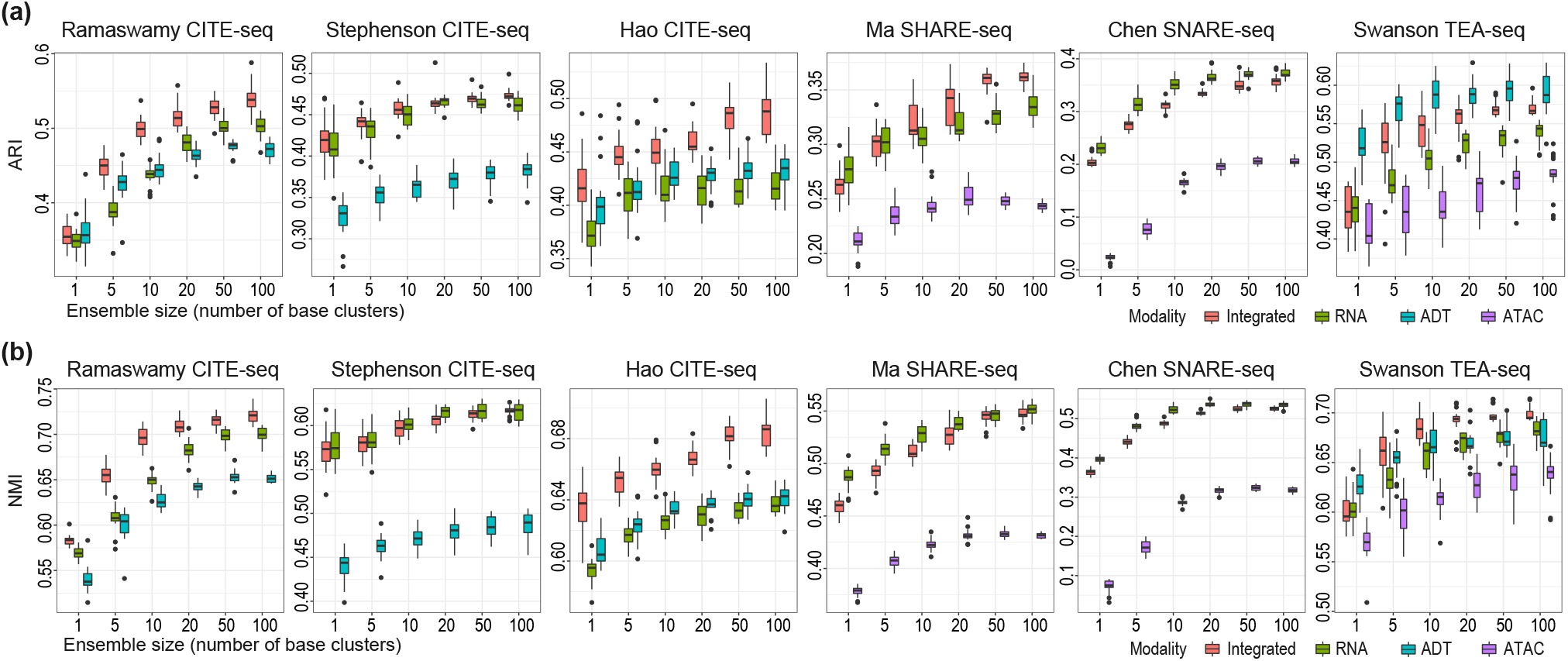
Comparison of integrated embedding of multimodality learned with those from unimodality using the k-means clustering algorithms. (**a**) Concordance of cell type clustering quantified by ARI on six multimodal single-cell omics data using SnapCCESS generated embeddings from either all modalities in a dataset or each data modality alone. (**b**) Similar to (a) but with the clustering concordance quantified by NMI. The entire procedure was repeated 20 times for capturing the performance variability.

### 3.3 SnapCCESS framework improves various clustering algorithms

Since the ensemble deep learning of embeddings in SnapCCESS is independent of clustering algorithms, we next tested the performance of SnapCCESS framework (epoch=1) by coupling it with a spectral clustering algorithm and SIMLR, a kernel-based clustering algorithm. Note that the two large CITE-seq datasets, Stephenson CITE-seq and Hao CITE-seq, were excluded due to the exponential growth of computational complexity with the number of cells in a dataset for the spectral clustering algorithm. Overall, we observed a clear increase in clustering performance with the increasing ensemble size, regardless of the types of clustering algorithms and concordance evaluation metrics (**Figure 5**). Nonetheless, compared to the simple k-means clustering algorithm, the application of more advanced SIMLR clustering and spectral clustering algorithms generally led to improved cell clustering as measured by their concordance to the cell type annotation. These findings are of particular interest to SIMLR, which was originally designed for analysing unimodal scRNA-seq data, as they demonstrate that embeddings learned by SnapCCESS from multimodal single-cell omics data can be used for a clustering algorithm designed for unimodal single-cell omics data.

**Figure 5.**
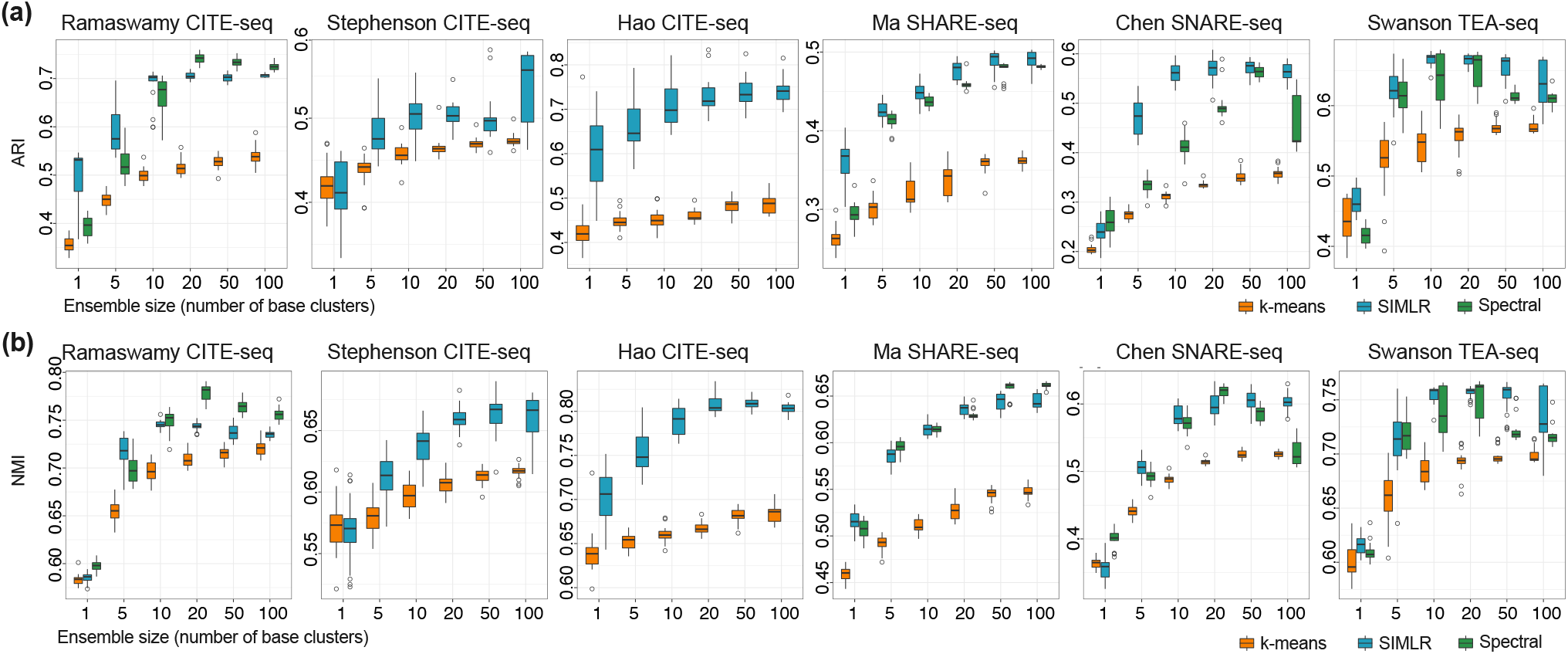
Evaluating the utility of the SnapCCESS framework with alternative clustering algorithms on multimodal single-cell omics data. (**a, b**) Concordance of the cell type annotations and cell clustering output from each clustering algorithm. The x-axis is the number of base clusters in the ensemble and the y-axis is the concordance quantification by either ARI (**a**) or NMI (**b**).

Similar to the results from k-means clustering, the improvement from SIMLR and spectral clustering also peaked around the ensemble size of 50 (**Figure 5**). This is also consistent with the results from scCCESS for scRNA-seq data analysis (Geddes *et al*., 2019), suggesting an ensemble size of 50 may be a suitable choice for the ensemble deep learning component in the SnapCCESS framework.

### 3.4 SnapCESS performs competitively to the state-of-the-art embedding generating methods for multimodal single-cell clustering

Various methods exist for generating embeddings from multiple data modalities in single-cell multimodal omics data. Some of the state-of-the-art examples include totalVI designed for combining RNA and ADT modalities in CITE-seq data and its sibling MultiVI for combining RNA and ATAC modalities such as in SHARE-seq and SNARE-seq, and Jvis-learn and MOFA+ which are generic and can be applied to all data modality combinations. Given the applicability of each method, we compared SnapCCESS (epoch=1 and ensemble size of 50) with TotalVI on the CITE-seq datasets, MultiVI on SHARE-seq and SNARE-seq datasets, and MOFA+ and Jvis-learn on all six datasets with two or three modalities. In particular, we used the multimodality integrated embeddings generated from each of these methods as input to k-means, SMILR clustering, and spectral clustering algorithms and examined the concordance of clustering output with cell type annotation using ARI (**Figure 6a**) and NMI (**Figure 6b**).

**Figure 6.**
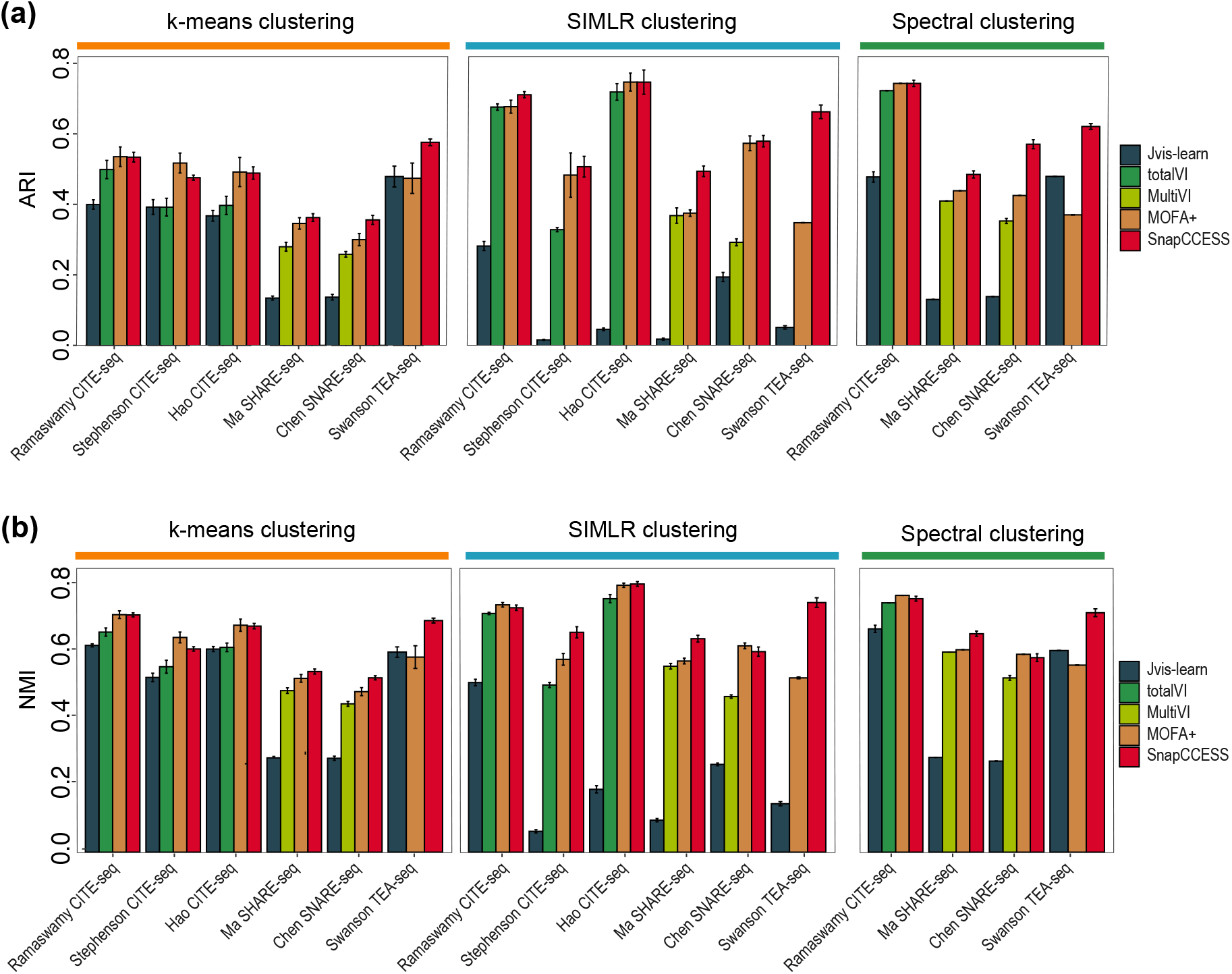
Comparison of SnapCCESS framework (epoch=1 and ensemble size of 50) with other multimodality embedding generation methods on cell clustering using k-means, SIMLR, and Spectral clustering algorithm. (**a**) Quantification of clustering concordance to cell type annotation using ARI. (**b**) Similar to (a) but quantifying clustering concordance using NMI. Each bar indicates the average performance across datasets with 20 repeats, and error bars represent the standard deviation.

We found that SnapCCESS performed competitively to other multimodality embedding generation methods. In most cases, its performance is significantly better than totalVI when applied to CITE-seq datasets and MultiVI when applied to SHARE-seq and SNARE-seq datasets (**Figure 6**). While MOFA+ also performed well especially on CITE-seq datasets, SnapCCESS appears to be slightly better than MOFA+ when used for generating embeddings and performing cell clustering on the SHARE-seq and the trimodal TEA-seq datasets. Interestingly, while clustering of the embeddings from Jvis-learn using k-means and Spectral clustering algorithms performed reasonably well, the use of SIMLR on Jvis-learn generated embeddings leads to poor results in many cases. These results may indicate the varying degree of generalisability of the multimodality embedding generation methods on different clustering algorithms. To this end, multimodality embeddings generated from the other four methods (i.e. SnapCCESS, totalVI, MultiVI, MOFA+) worked well regardless of the used clustering algorithm.

## 4 Discussion and conclusion

A main challenge in single-cell omics data analysis is in handling the high-dimensionality of the feature space (Yang *et al*., 2021). Embedding learning is a popular approach for reducing feature dimension for subsequent analysis such as clustering of cells. While many methods have been designed to generate embeddings for unimodality scRNA-seq data, such methods could not be directly applied for integrating multiple data modalities in multimodal single-cell omics data. The development of methods that can learn integrated embeddings across multiple data modalities, while still in its infancy, is essential for the dimension reduction of multimodal single-cell omics data.

A key innovation in SnapCCESS is the adaptation of the snapshot ensemble learning technique (Huang *et al*., 2017) which significantly reduce the computation time and resource for generating multi-view of embeddings compared to conventional VAE ensembles. Nevertheless, the clustering of each embedding generated from SnapCCESS is still performed individually. While parallelisation can be implemented to speed up the process, designing clustering algorithms that can cluster cells via multiple embeddings simultaneously could further improve computational efficieny. Related to this, there are various ensemble deep learning methods that aim to reduce computation time by using techniques such as model branching and neuron deactivation (Cao *et al*., 2020). The effectiveness of these alternative ensemble deep learning approaches for learning embeddings from multimodal single-cell omics data remains to be tested.

The utility of the embeddings generated from multimodal single-cell omics data is much wider than cell clustering. While clustering is one application that can make use of embeddings learned from such data, other tasks such as supervised cell type classification (Abdelaal *et al*., 2019) and unsupervised number of cell type estimation (Yu *et al*., 2022) that take the embeddings as input can also be applied for analysing multimodal single-cell omics data. Therefore, designing methods that can generate better embeddings will impact various downstream analyses and applications of multimodal single-cell omics data. To this end, the utility of ensemble deep learning methods for these applications (e.g., cell type classification, number of cell type estimation) should be investigated in future studies.

While the current study evaluates SnapCCESS framework for clustering cells into discrete groups, the cell type structures from many biological systems are hierarchical, with subpopulations of cells existing in each major cell type (Wu and Wu, 2020). The development of multimodal single-cell omics technologies facilitates the characterisation of such hierarchical cell type relationships. Therefore, developing methods that are capable of multi-resolution clustering of cells on datasets with multimodal molecular attributes is a direction of future research. Another recent expansion in the single-cell omics field is the increasing availability of spatial single-cell omics data produced by an array of new spatial profiling technologies (Larsson *et al*., 2021). Combining spatial data with other omics data types produced from the same cells and samples has the potential to uncover a wealth of information, including spatial-related cell type structure, and will help us gain a deeper understanding of cell and tissue development and disease progression. Developing clustering algorithms for integrating spatial data with other omics datasets is challenging and requires further methodological innovation.

In summary, our previous work demonstrated that ensemble learning of embeddings provides an effective approach for improving downstream clustering analyses by providing a multi-view of the input data (Geddes *et al*., 2019). Here we extend this idea for single-cell multimodal omics data analysis by introducing SnapCCESS, an efficient ensemble deep learning framework using VAE and snapshot techniques, gaining high performance in cell clustering while alleviating the limitation on computation efficiency in conventional ensemble learning methods. Since the clustering of individual embeddings can be performed independently from each other, the proposed framework can benefit from further speed up by parallelisation of embedding clustering. We expect SnapCCESS to serve as a useful tool and spark the future development of ensemble deep learning methods for multimodal single-cell omics data analysis.

## Acknowledgments

The authors would like to thank all their colleagues, particularly those at the Sydney Precision Data Science Centre, The University of Sydney, for their intellectual engagement and constructive feedback.

## Author contributions

P.Y. conceived the study. L.Y. led the data analysis with input from P.Y., C.L. and J.Y.H.Y.; L.Y. and P.Y. wrote the manuscript with input from C.L. and J.Y.H.Y.; All authors read and approved the final manuscript.

## Data availability

All data used in this study are publicly available. Details of each dataset are reported in Section 2. The accession links are summarised here including the *Ramaswamy CITE-seq dataset* (GSE166489), *Stephenson CITE-seq dataset* (E-MTAB-10026), *Hao CITE-seq dataset* (GSE164378), *Chen SNARE-seq dataset* (GSE126074), *Ma SHARE-seq dataset* (GSE140203), *Swanson TEA-seq* (GSE158013).

## Code availability

SnapCCESS is implemented as a Python package and is freely available from https://github.com/yulijia/SnapCCESS.

